# Human Embryonic Expression Identifies Novel Essential Gene Candidates

**DOI:** 10.1101/2020.08.15.252338

**Authors:** Monica Penon-Portmann, Jiyoo Chang, David R. Blair, Beatriz Rodriguez-Alonso, Hakan Cakmak, Aleksandar Rajkovic, Joseph T. Shieh

## Abstract

Disruption of essential genes leads to pregnancy loss, early lethality, or severe disease. Current methods to predict genes that underlie severe phenotypes include knockout animal model systems, evolutionary conservation, and variation intolerance metrics. With existing methods, human lethal genes are missed due to interspecies differences or paucity of gene characterization. We analyzed global gene expression in stages of early human development (1-cell to the blastocyst). These data were integrated with all 4049 current murine knockout phenotypes, genome-wide evolutionary gene conservation, and human genic intolerance metrics. We found that currently established human essential genes and orthologs of murine essential genes demonstrate higher gene expression across developmental stages compared to non-essential genes (Wilcoxon rank sum test, p<8.5e-10), indicating that higher expression correlates with essentiality. Of 1438 unique genes candidates with the highest expression, an estimated 1115 (78%) have not yet been associated with human disease and are thus novel candidates. The essential gene candidates concur with four prediction metrics, further supporting essentiality. We also assessed gene-specific expression changes during early development for their ability to predict essentiality. Genes that increase in expression were more likely to be essential (Fishers exact test, p<2.4e-06), suggesting that dynamic temporal expression during development may be particularly important. We find that embryonic gene expression can be used to prioritize genes that currently lack a Mendelian phenotype. Human embryonic gene expression is readily available, and applied as a novel tool, it may identify highly conserved processes vital in development.

## Introduction

Essential genes are critical for the survival of an organism (Bartha, Di Iulio, Venter, & Telenti, 2018). Their disruption may lead to pregnancy loss, early lethality or severe disease. By identifying genes that are essential for early human development, we can provide insights into the molecular basis of fitness and reproductive success (Ortega, Winblad, Plaza Reyes, & Lanner, 2018; T. Wang et al., 2015). While high-throughput sequencing technologies facilitate the detection and interpretation of variants in well-annotated genes, they also generate thousands of candidates that can only be interpreted in light of our current knowledge. Even with an increasing insight into molecular pathophysiology, gene-disease annotation covers only about 25% of genes in the genome, leaving thousands with no known human phenotype correlation (Amberger, Bocchini, Scott, & Hamosh, 2019). Furthermore, despite advances in diagnostics, poor pregnancy outcomes remain unexplained in at least 30% of cases (Rajcan-Separovic et al., 2010; Ray, Shah, Gudi, & Homburg, 2012; Yatsenko & Rajkovic, 2019). To facilitate a broader discovery process, it is critical to have tools that can assist in the prediction of essential genes which in turn may help interpretation of novel variants that arise with genomic data.

Current methods to predict essential genes that underlie severe phenotypes include knockout animal model systems, evolutionary gene conservation, gene disruption in haploid cell lines, and the study of human development through sequencing (Bartha et al., 2018; Collins et al., 2019; Fogarty et al., 2017; Fraser, 2015; Muñoz-Fuentes et al., 2018; Petropoulos et al., 2016; Qiao et al., 2016; Suzuki et al., 2018; T. Wang et al., 2015). Model systems allow for the observation of shared phenotypic traits between species (Mee et al., 2005; Ortega et al., 2018). This approach has successfully identified genes that are critical in early murine development, which appear to translate well to humans (Mee et al., 2005; Muñoz-Fuentes et al., 2018). For instance, homozygous pathogenic variants in *HYLS1* lead to perinatal human and mouse lethality (OMIM #236680) (Mee et al., 2005). Even though studies of mouse embryo development have provided key insights into early developmental pathways, species-specific differences limit the extrapolation of some findings to humans. Translation is challenging in many instances due to inter-species differences such as those observed in the timing and mechanisms of lineage specification (Ortega et al., 2018; Xie et al., 2010). For instance, humans begin specification of the trophectoderm and inner cell mass in the blastocyst stage while, in contrast, mice begin this differentiation earlier in the morula stage (Niakan & Eggan, 2013; Ortega et al., 2018). Such differences in the timing and in the sequence of events between species elucidate the importance of human-specific analyses for the prediction of essential genes.

Genomic data from large populations has also served to predict essential genes (Karczewski et al., 2020; Lek et al., 2016). For example, multiple statistical methods have been developed to estimate the extent to which a particular genetic locus is intolerant to different types of sequence variation (Ge et al., 2016; Ge, Kwok, & Shieh, 2015; Havrilla, Pedersen, Layer, & Quinlan, 2019; Karczewski et al., 2020; Lek et al., 2016). Regions of the genome highlighted by these measurements may be under evolutionary constraint due to their effects on fitness, including embryonic lethality. That said, not all essential genes (for example, those underlying recessive conditions) show obvious patterns of such constraint (Dawes, Lek, & Cooper, 2019), necessitating the development of orthogonal methods using additional datasets to further increase our knowledge of human essential genes.

Additional approaches for essential human gene identification and characterization include human-specific analyses such as genomic sequencing of individuals with recurrent pregnancy loss (Qiao et al., 2016; Robbins, Thimm, Valle, & Jelin, 2019; Suzuki et al., 2018; Yatsenko & Rajkovic, 2019), single-cell RNA sequencing in early human development (Petropoulos et al., 2016) and genome editing (Fogarty et al., 2017). These methods have identified other critical genes; nonetheless, many candidate genes remain undiscovered in terms of their association to human fitness.

We hypothesized that gene expression patterns during early human development can predict essential genes and identify novel candidates. To test our hypothesis, we analyzed global gene expression in all stages of early human development (1-cell to the blastocyst). We first assessed potential novel human essential genes by comparing orthologous human genes to those knocked-out in the mice and to those highly conserved down to drosophila. We then compared gene expression to metrics that predict essential genes including all current murine knockout phenotypes, genome-wide evolutionary gene conservation, and human genic intolerance. We test our predictions by comparing attributes across essential and non-essential genes. Our analyses suggest that human gene expression during development is a useful tool for essential gene prioritization. Furthermore, we provide a list of novel gene candidates based on expression data and other predictive attributes.

## Methods

### Murine knockout phenotype data

We extracted mouse knockout phenotype data from the International Mouse Phenotyping Consortium (IMPC), release 9.2 (ftp://ftp.ebi.ac.uk/pub/databases/impc/release-9.2/csv/, access date: March 9^th^, 2020). From this dataset, we identified all homozygous or hemizygous knockdown alleles that lead to lethality and are therefore essential in the mouse. We defined lethality using the following mouse phenotype terms: *embryonic lethality prior to organogenesis*, *embryonic lethality prior to tooth bud stage*, *prenatal lethality*, *prenatal lethality prior to heart atrial septation*, *preweaning lethality complete penetrance*, and *preweaning lethality incomplete penetrance*. Each gene was considered essential if it had at least one lethal phenotype. We included all data provided by the IMPC resource and recovered a total of 4049 unique murine knockout (KO) genes.

### Gene orthologs for mouse and drosophila melanogaster

We used Ensembl BioMart to extract human gene orthologs to *Mus musculus* and *Drosophila melanogaster* (access date: March 10^th^, 2020). We queried the human genes dataset from GRCh38. We compared various attributes across orthologous genes sets including transcript, homology type, and percent sequence identity. We included genes with one-to-one orthologs and excluded those with multiple orthologs to minimize the chances of misalignment.

### Human constraint data

We used human constraint data from gnomAD version 2.1.1 (access date: March 10^th^, 2020). We identified constraint scores per gene for all human genes, which were then used for downstream analyses (n=19,689 genes with constraint scores). Specifically, we used constraint metrics that represent the intolerance to loss-of-function variation (pLI score) and the intolerance to missense variation (missense Z-score). The pLI score represents the probability that a gene is loss-of-function (LoF) intolerant, with the most intolerant genes having a pLI ≥ 0.9 while the most tolerant genes possessing a pLI ≤ 0.1 (Lek et al., 2016). We also looked at the missense Z-score, which is a gene level estimation of intolerance to missense variants. Thus, genes with high Z-scores are intolerant to missense variation (Lek et al., 2016). Additionally, we used the ratio of observed to expected variation (OE ratio) to measure genic intolerance to variation. Genes that have a low OE ratio are also predicted to be intolerant to variation and potentially subject to negative selection (Karczewski et al., 2020; Lek et al., 2016). We also extracted a list of 1815 genes tolerant to biallelic loss-of-function variation (Karczewski et al., 2020).

### Cell essential genes

A dataset of cell essential genes was extracted from a genome-wide study that screened for essentiality in haploid human cell lines (Hart et al., 2015). We defined cell essential genes as those showing a fitness defect in all five cell lines (n = 829), which are presumably less likely to be context-dependent essential genes.

### Expression data

We obtained human developmental expression data from the NCBI Gene Expression Omnibus (http://www.ncbi.nlm.nih.gov/geo, under accession GSE18290). The data set contained array expression profiles obtained with the Affymetrix Human Genome U133 Plus 2.0 Array and included expression from human embryonic development at the 1-cell, 2-cell, 4-cell, 8-cell, 16-cell or morula, and 32-cell or blastocyst stages. We verified that the expression data was normalized. This dataset contained a total of 54,676 probes, representing approximately 22,190 genes. Expression of the three replicates in each probe was averaged to generate data representing each embryonic cell stage. The corresponding gene symbols were appended to each probe set. To analyze expression, we determined the change in expression between stages 1-cell to 4-cell, which include transcripts of maternal origin, in addition to later stages, which include genes expressed only by the zygote.

We obtained tissue expression data from the NIH Genotype-Tissue Expression (GTEx) (V6, October 2015) project through the table browser feature on UCSC Genome Browser (http://genome.ucsc.edu/, access date: May 10^th^, 2020). We also extracted single-cell RNA-seq expression data from Petropoulos et al. to identify the lineage with the highest ranked expression per gene (Petropoulos et al., 2016, access date: May 1^st^, 2020). Each gene was appended the lineage with their highest ranked expression during embryo day 5 – 7. We also estimated lineage representation for the genes within the 95^th^ percentile of embryonic expression. Finally, enrichment analyses of developmental genes were performed using the online David bioinformatics database (https://david.ncifcrf.gov/).

### Human phenotype data and known human lethal genes

All currently annotated human phenotype genes were downloaded from Online Mendelian Inheritance in Man (OMIM, https://omim.org/downloads/, access date: November 21^st^, 2019). Furthermore, to identify those genes known to cause prenatal or perinatal lethality in humans, we used all that contained terms associated with lethality before or shortly after birth in the clinical description using information from Dawes *et al.*, 2019 (n = 624 genes). Dataset alignment was completed using gene IDs, and canonical isoforms were used in instances with multiple isoforms.

### Statistical Analyses

We performed statistical tests, data analyses and visualization with R software, version 1.2.1335. Figures were made with R graphics and the ggplot2 package. The Wilcoxon rank sum test was used to compare the distributions of constraint scores and genic expression between essential and non-essential genes. We measured the association between attributes such as nucleotide identity or embryonic expression and genic constraint using Spearman’s rank correlation. We estimated an odds ratio to determine essential gene enrichment among genes with up-trending versus down-trending expression. A Fisher’s exact test of independence was used to compare predictive traits across groups of genes. For gene function enrichment analyses, multiple testing correction was performing using the Bonferroni and Benjamini-Hochberg methods (https://david.ncifcrf.gov/).

## Results

We identified a total of 4049 unique murine genes with homozygous or hemizygous knockout models (Table 1). From the 4049 genes, 1705 lead to a lethal phenotype in mice. Within this group of murine essential genes, 439 (26%) lead to prenatal mortality and 1266 (74%) to postnatal mortality. Only four genes were lethal in the hemizygous state (*Phf6, Prps1, Kdm5c, Fgd1*). Each gene was then annotated to its human ortholog. From 4049 unique murine knockout genes, 3580 (88%) had one-to-one human orthologs. Out of the 3580 orthologs, 648 (41%) genes were annotated in OMIM with a known human phenotype. This initial analysis identified that many essential murine genes do not have a known human phenotype to date, suggesting a potential gap in human phenotypic annotation or inter-species correlation.

**Table 1.** Human orthologs of murine essential and non-essential genes, including the subset with human phenotype annotation.

| **Murine Phenotype** | **Murine KO genes** | **Human Orthologs** | **Genes with a known human phenotype** |
| --- | --- | --- | --- |
| **Essential** | 1705 | 1578 | 648 (41%) |
| Prenatal mortality | 439 | 400 | 160 |
| Postnatal mortality | 1266 | 1178 | 488 |
| **Non-essential** | 2344 | 2002 | 465 (23%) |
| Phenotype | 2336 | 1994 | 461 |
| No phenotype | 8 | 8 | 4 |
| **Total** | 4049 | 3580 | 1113 |

### Murine knockout models predict essential human genes

We next tested whether human gene intolerance scores differ between essential and non-essential genes. The 3580 human orthologs with current murine knockout phenotypes, corresponding to all of the knocked-out genes in mice were merged with constraint data derived from human population sequencing (Figure 1). Human orthologs with constraint scores were divided into murine essential (n = 1515) and murine non-essential genes (n = 1896) depending on whether their knock-out led to model lethality. We then used population loss-of-function and missense variant constraint metrics to determine whether essential and non-essential genes differ in their tolerance to variation. For these constraint measurements, a low observed to expected ratio (OE ratio) indicates that a gene is likely to be more intolerant to genetic variation. The distribution of constraint scores between known essential and non-essential human genes yielded significant differences between the groups (Figure 1). Notably, human orthologs of murine essential genes have less variation, or a lower OE ratio, compared to murine non-essential genes. Constraint was significantly greater in murine essential genes for both loss-of-function (LoF) and missense variation (Wilcoxon Rank Sum test, p < 2.2e-16,). Hence, human orthologs with a mouse model leading to lethality are more intolerant to variation compared to non-lethal genes.

**Figure 1.**
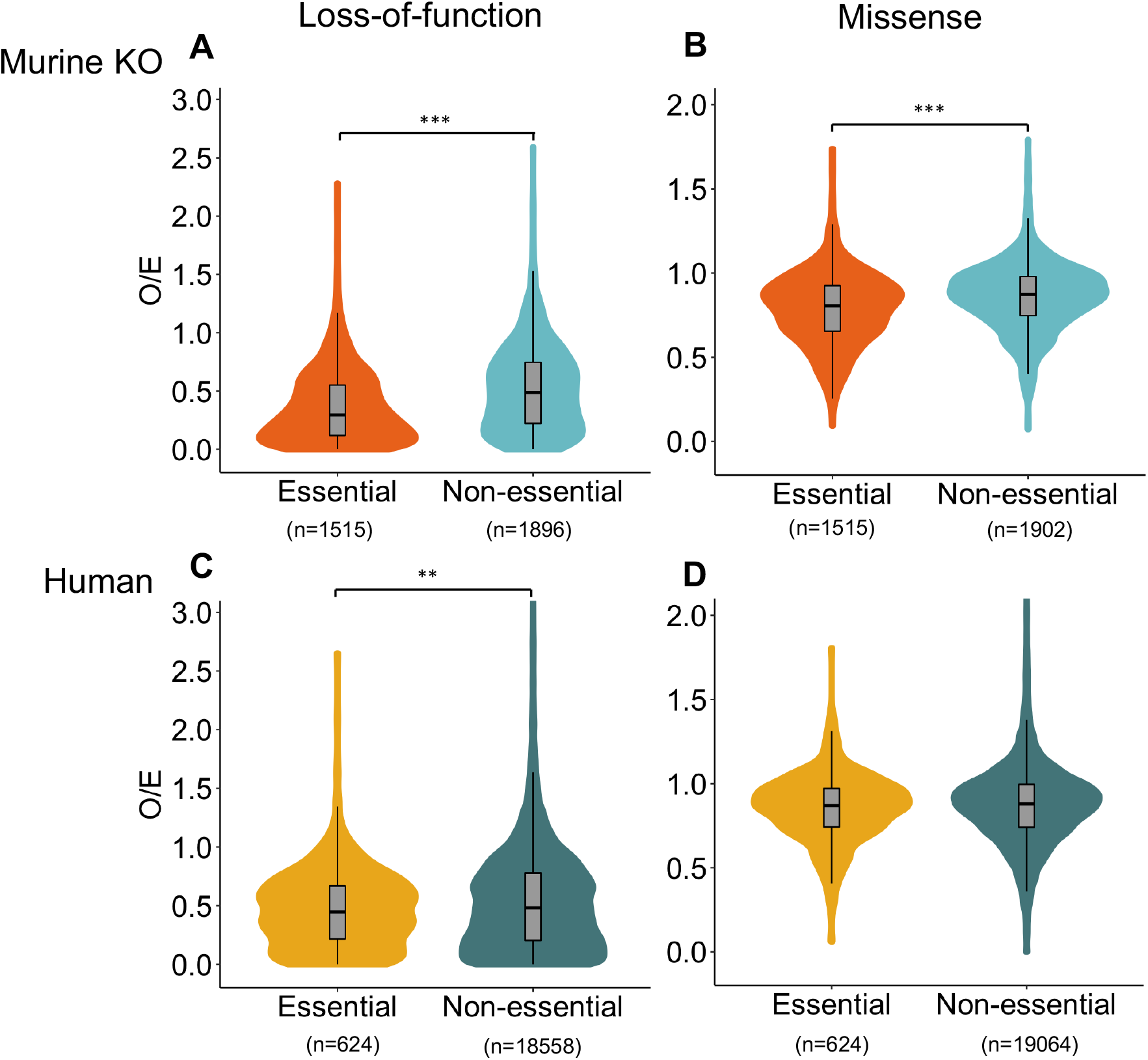
Human constraint correlates with essentiality. A) Comparison of human loss of function (LoF) variant observed-to-expected (OE) ratios between human orthologs of murine essential genes and non-essential murine orthologs (Wilcoxon rank sum test, p<2.2e-16). Genome-wide knockout data represented in graph. B) Comparison of the OE ratio for missense variation between human orthologs of murine essential genes and non-essential murine orthologs (Wilcoxon rank sum test, p<2.2e-16). C) Comparison of the LoF OE ratio between human essential genes and non-essential genes (Wilcoxon rank sum test, p<0.007). D) Comparison of the missense OE ratio between human essential genes and non-essential genes (Wilcoxon rank sum test, p=0.5428).

To help validate this finding, we also examined human essential genes that lead to a lethal human phenotype (n=624) and found that these also demonstrated LoF constraint when compared to non-essential genes (Figure 1) (Wilcoxon rank sum test, p<0.007). However, known human essential genes were not significantly constrained with respect to missense variation (Wilcoxon rank sum test, p = 0.5), suggesting no detectable effect of missense variants on this particular feature. Human constraint therefore seems to have some correlation with lethality; however, this prediction method is not without limitation.

Remarkably, we found 842 (56%) human orthologs of murine essential genes do not show evidence for LoF constraint in humans (pLI ≤ 0.2). Moreover, the opposite is also true, some constrained genes have a non-lethal mouse phenotype. Constraint metrics vary depending on the mode of phenotype inheritance, for instance, genes with recessive inheritance, may be more tolerant to variation. Thus, essential genes that exhibit haplosufficiency may be missed by constraint metrics.

### Interspecies conservation predicts essential human genes

Since some genes involved in early human development are predicted to be highly conserved across species, we also assessed conservation between human and drosophila. A genome-wide query identified a total of 3082 one-to-one orthologs between humans and *Drosophila melanogaster,* which vary in their degree of interspecies conservation. We compared nucleotide similarity to human population-based genic constraint. We hypothesized that highly conserved genes across species should have a critical function and therefore are subject to strong negative selection if mutated. We found a positive correlation between interspecies nucleotide sequence identity and human population genetic constraint (Spearman’s rank correlation, *R* = 0.27, p < 2.2e-16) (Figure 2). Consequently, genes with higher nucleotide similarity were more likely to be intolerant to variation. Haploinsufficient genes in humans are those that are intolerant to LoF variation and therefore have a high pLI score. Genes with high pLI scores also tend to share a greater interspecies nucleotide sequence similarity, as they clustered in the right upper quadrant of the scatter plot (Figure 2A). This suggests that genes that have a close interspecies resemblance are less likely to show LoF variation in the human population.

**Figure 2.**
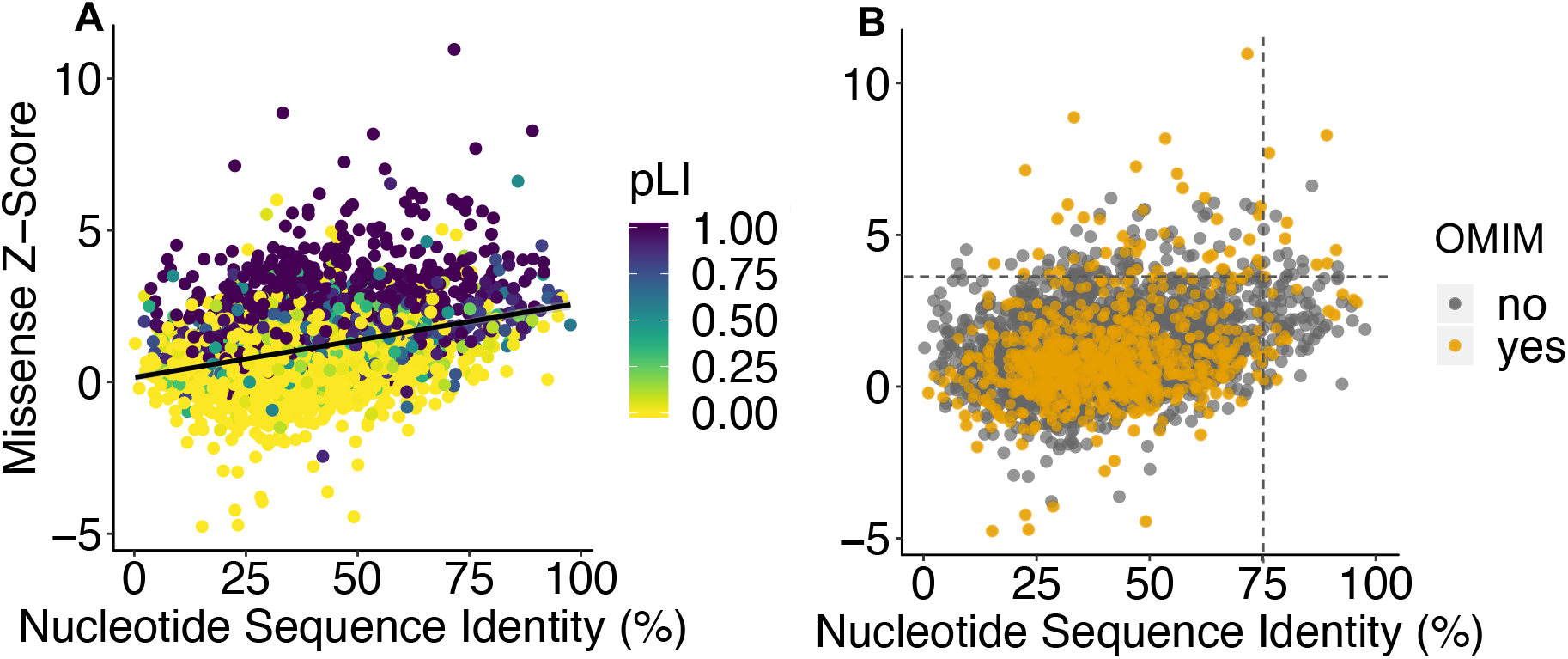
Gene conservation correlates with constraint. A) Nucleotide sequence identity of one-to-one orthologs between *Drosophila melanogaster* and *Homo sapiens* compared to the population missense constraint (Z-score). Each dot represents an ortholog (n = 2987 genes). There is a positive association between conservation and constraint (Spearman’s rank correlation, *R* = 0.27, p < 2.2e-16) B) Same data as A colored by genic representation in OMIM (orange). The top right-hand quadrant delimits genes in the top 5% of missense variation depletion (Z-score ≥ 3.9) and nucleotide identity conservation (≥75%).

We also determined how many of these cross-species orthologs between humans and drosophila have a known human disease reported in OMIM (Figure 2B). From a total of 3082 orthologs, 931 (30%) have a known human phenotype, yet a striking 2151 (70%) remain unannotated. We identified a subset of 16 orthologs with nucleotide sequence identity ≥ 75% and high constraint (missense Z-score ≥ 3.9) that could be studied in the future. We propose that these genes might make interesting candidates as potential essential genes given their attributes. Nine genes from 16 candidates have not been associated with any known human disease at the time this manuscript was written (Table S1). Remarkably, although conservation may serve to identify some genes with strong negative selection pressure, some genes with high intolerance scores in humans have low nucleotide resemblance to drosophila, reflecting a possible divergence in biological function across species. Similarly, from the 624 known essential human genes, 181 had a mouse knockout model, and 39 of these 181 (21.5%) genes were non-essential in mice yet essential in human suggesting further human studies are needed. Altogether, our multi-parameter analysis of essentiality metrics demonstrates that constraint, conservation and murine model lethality correlate with essentiality and identified potential novel candidates.

Nonetheless, these metrics showed some limitations in translation likely due to interspecies differences.

### Essential gene expression in early human development

Given its potential impact on early human development, we examined whether gene expression during human development predicts essentiality. We hypothesized that gene expression patterns during early development can predict essential genes and identify new candidates for lethality. To test our hypothesis, we analyzed genome-wide gene expression in all stages of early human development (1-cell to the blastocyst) (Xie et al., 2010). We compared levels of expression of human essential and human orthologs of murine essential genes to their non-essential counterparts (Figure 3). We found that established human essential genes and orthologs of murine essential genes demonstrated higher gene expression in all developmental stages compared to non-essential genes, indicating that higher expression in early development correlates with essentiality (Figure 3, Wilcoxon rank sum test, p<8.5e-10). Furthermore, when we randomized genes in multiple permutations, this difference was lost, suggesting this relationship was unlikely to be due to chance.

**Figure 3.**
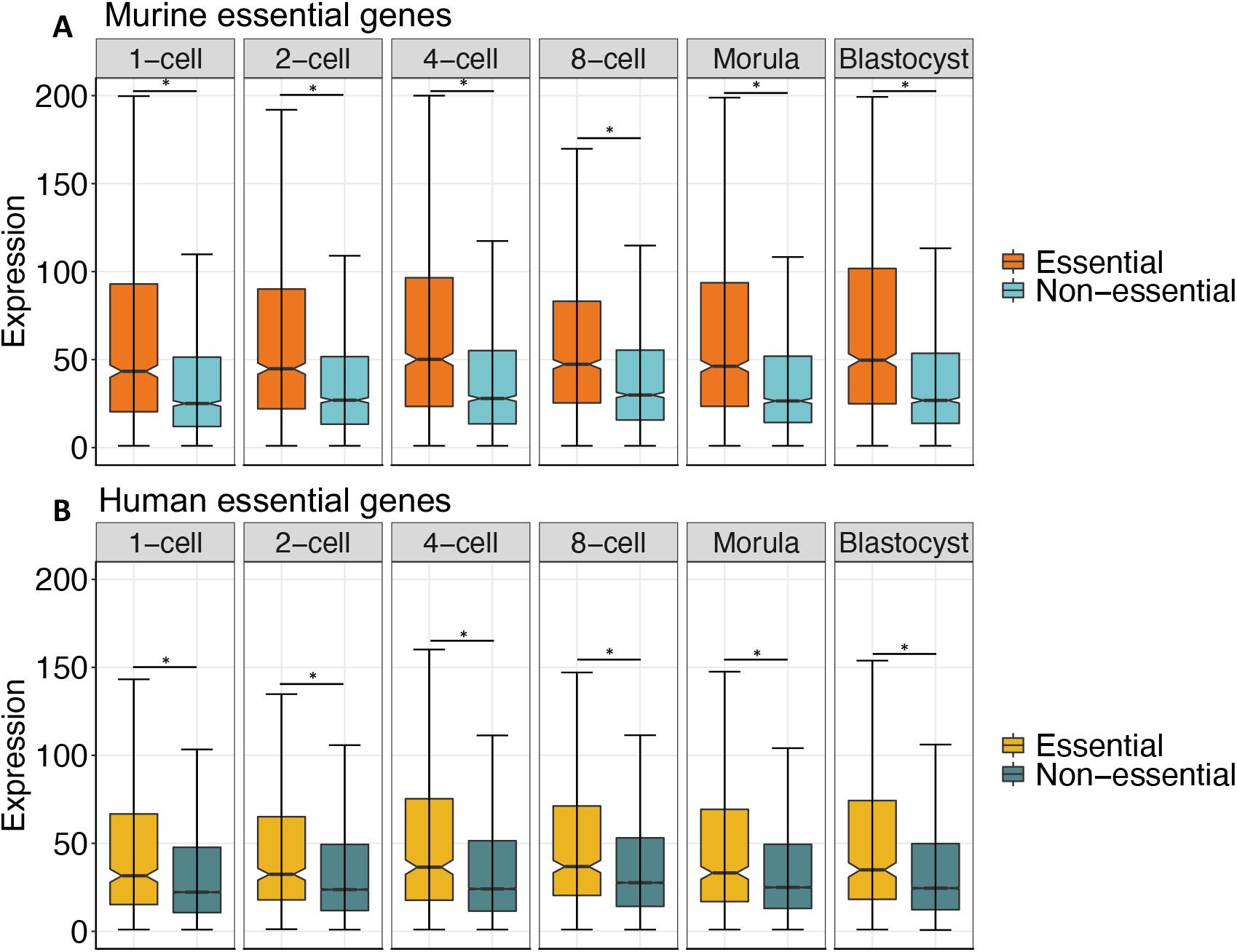
Gene expression and essential genes. High expression in early human development is associated with human and murine essentiality. A) Human orthologs of murine essential genes (n=1504 in orange) that lead to mouse lethality are more highly expressed in early human stages compared to non-essential orthologs (n=1853 in light blue). B) Known human essential genes (n=604 in golden yellow) have higher expression compared to non-essential human genes (n= 21,587 in dark blue). These differences were significant in all developmental stages (Wilcoxon rank rum test, *p<8.5e-10), whereas expression of randomized genes was not.

We then assessed for association between developmental expression levels and other attributes such as conservation and genomewide constraint. Indeed, highly expressed genes tended to be evolutionarily conserved, and those transcribed after zygote genome activation (cell stages 8-cell to blastocyst) demonstrated more intolerance to variation. The correlation with constraint was observed for both for both loss-of-function and missense variation; however, this association was absent in earlier cell stages when gene expression is mostly driven by transcripts of maternal origin.

We identified the embryonic lineage for genes in the top 95^th^ percentile with respect to embryonic expression (Table S2) using data from Petropoulous *et al.* (Petropoulos et al., 2016). Approximately 25% of highly expressed genes were primarily expressed in the epiblast, which gives rise to the embryo. The remaining 75% of genes are expressed in cells that go on to make the trophectoderm for the placenta and the primitive endoderm for the yolk sac/visceral endoderm. Hence, highly expressed genes in early development were expressed across the three cell types suggesting potentially important roles in each developing structure.

### Patterns in gene expression correlate with essentiality

Since stage-specific gene expression analyses correlated with essentiality, we next assessed trends in expression. The most upregulated genes (Figure 4) showed an enrichment for human disease: 16 of 36 genes (or 44%) as they were annotated as disease-associated in OMIM. In addition, several known essential human genes were found among this set, including *NLRP7*, which is a known cause of recurrent pregnancy loss and hydatidiform mole (Aghajanova et al., 2015). Other genes in this group were noted to cause lethality at an early age such as *NDUFS6* (mitochondrial complex I deficiency, nuclear type 9) and *HMOX1* (Heme oxygenase-1 deficiency). Another gene identified in this set is *LIN28A*, which is not currently known to be associated with a human phenotype but is one of the factors, along with *OCT4, SOX2* and *NANOG*, that can reprogram somatic cells to pluripotent stem cells (Yu et al., 2007).

**Figure 4.**
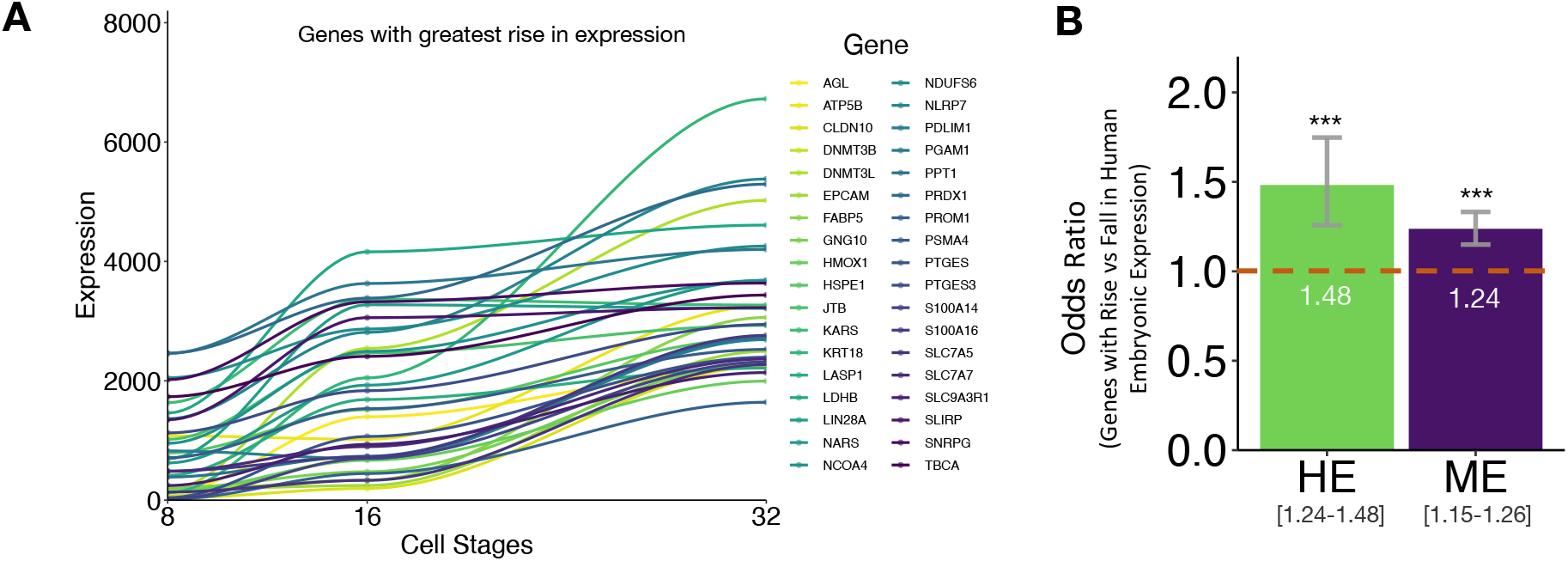
Up-trending zygote-activated genes correlate to essentiality. A) Top ranked upregulated genes between the 8-cell and 32-cell stage (blastocyst). B) Odds ratio (OR) analyses comparing the number of genes that increase to those that fall in expression. Zygote-activated up-regulated genes are significantly more likely to be essential (HE = human essential) (ME = murine essential orthologs) (OR = 1.48 for HE and OR= 1.24 for HE). Brackets represent the 95% confidence interval, orange hatched line drawn at OR=1. (Fisher’s exact test, ***p<2.36e-06).

**Figure 5.**
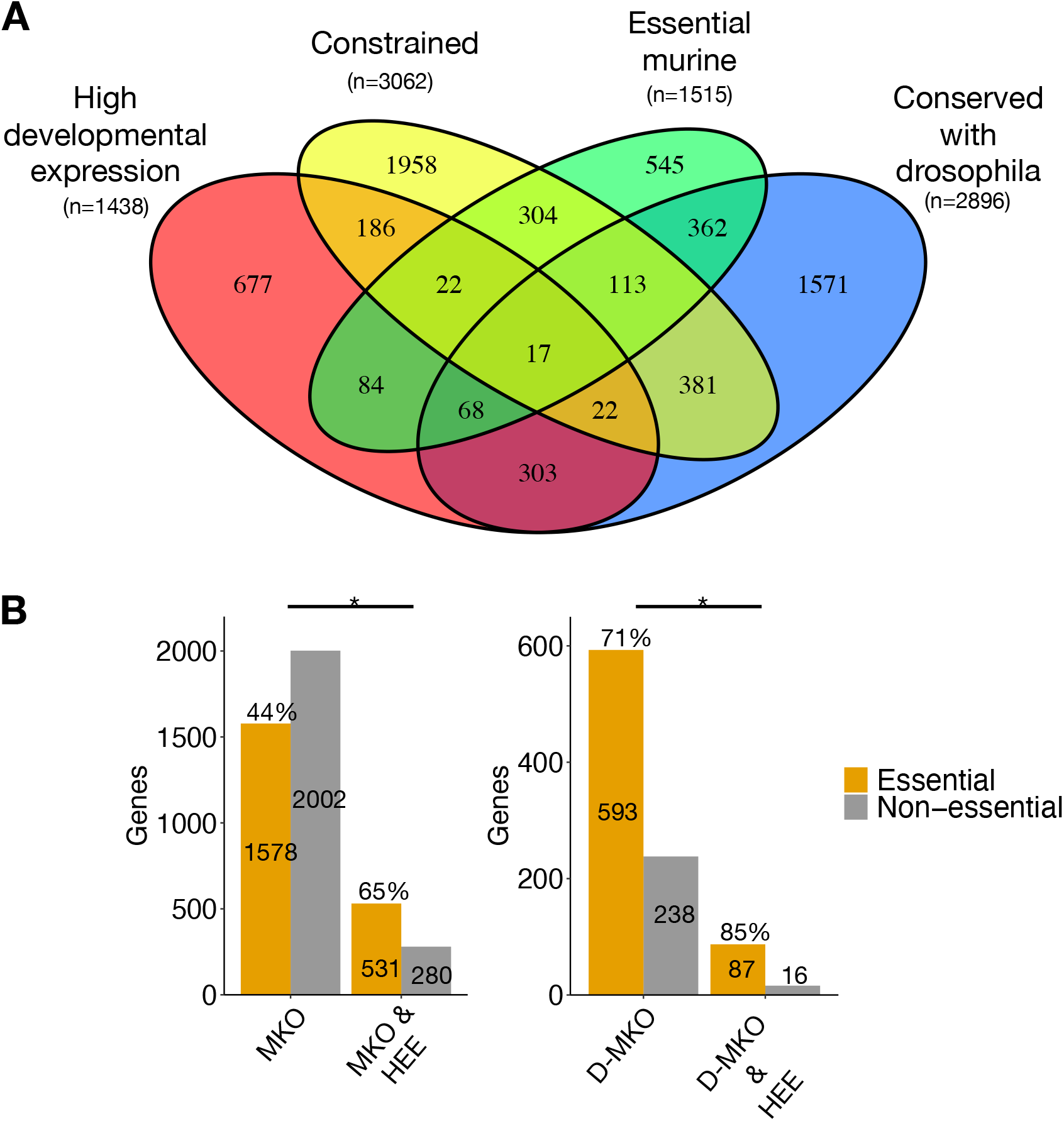
Genes with overlapping predictive traits are enriched for essentiality. A) Venn diagram illustrating the number of genes within the intersection of traits that relate to essentiality. B) Frequency of genes with a mouse knockout model (MKO, left) and MKO with a drosophila ortholog (D-MKO, right), thus highly conserved. There is a significantly higher fraction of essential genes when these genes are also highly expressed in development (HEE, Fisher’s exact test, p<0.05).

Accordingly, other genes with a rise in expression in these early stages could be interesting essential gene candidates. In zygote-activated genes (cell stages 8-32), the change in expression ranged from 0.03 to 54-fold (Figure 4A). To systematically assess for enrichment of essentiality in up-or down-trending zygote activated genes, we computed an odds ratio for essentiality among genes that rise versus those that fall in expression. Not surprisingly, genes with a rise in expression were more likely to be essential when compared to those that decreased (Figure 4B, Fisher’s exact test, ***p<2.373e-06). Depending on their underlying function, we surmised that genes with high expression in development may either fluctuate in time or remain highly expressed throughout the lifetime of an organism. Therefore, we searched for genes that displayed temporal differences in expression between early development and postnatally. We identified a number of genes whose expression level decreased significantly from the top 10% in early development to the bottom 10% rank in the adult. We found 66 unique genes that were highly expressed early in development but decreased postnatally. A systematic comparison of the human orthologs for murine essential genes demonstrated that expression of essential genes remained significantly higher than non-essential genes postnatally, suggesting some of these genes may exhibit ongoing functionality. To test the functional role of decreasing genes versus those that remained highly expressed, we performed gene set enrichment analyses of both groups. Ontology analyses of the most highly expressed genes in the early embryo (95^th^ percentile) demonstrated that these candidates showed pathway enrichment in ribosomal units, translation and metabolic enzymes, suggesting these processes might be important for future consideration in essentiality. Analyses of the 66 gene candidates with temporal high expression prenatally yielded a significant enrichment in apoptosis-related proteins and DNA binding.

### Genes that share multiple attributes are enriched for essentiality

To consider how the attributes might capture essential genes collectively, we compared the gene candidate lists generated from the following attributes: conservation, constraint, high embryonic expression and model organism lethality. Most genes of these were highlighted by multiple attributes, and a group of 17 were in the gene lists for all of them (Figure 6A).

We also noted that genes that shared attributes were significantly enriched for essentiality (Figure 6B, p<0.05 Fisher’s exact test). In addition, we assessed 1438 unique genes candidates with the highest expression (95^th^ percentile), and we estimated that 1115 (78%) have not yet been associated with human disease and are thus novel candidates. We also verified that only 50 genes from a list of 1812 (3%) are known to tolerate biallelic LoF in humans and were thus removed from our list of candidates. Furthermore, the majority of gene candidates with the highest expression in early development were also highlighted by our other predictive metrics, further supporting their essentiality. We also determined if these novel candidates overlapped with proposed essential genes from cell viability (Hart et al., 2015) and found that 629 of our highly expressed genes are absent from the predicted human cell essential genes.

## Discussion

In recent years, researchers have elucidated many elements of human preimplantation biology and essential genes, but many of these genes likely remain uncharacterized. In this study, we systematically assessed the attributes of essential human genes and demonstrate that developmental expression is a novel characteristic that can be used predict and prioritize essential gene candidates. In the future, high throughput genetic sequencing for embryonic arrest, pregnancy loss and infertility will likely identify many additional variants, which indicates a need for tools to prioritize candidates and facilitate the interpretation of such variants with uncertain significance.

Current tools to predict human gene essentiality include model systems, gene conservation and population constraint data. However, human lethality could be missed by current prediction tools when there is a lack of a human phenotype, if there are differences between the model organism and the human, and when genes lack annotation. Thus, human genic expression may complement prediction tools and may preclude the need for translation across systems.

Our study demonstrates how several metrics help predict essential genes but are subject to the challenges of translation. For example, mouse lethality is a good predictor of human lethality. In fact, approximately 75% of known human prenatal-lethal genes are associated with knockout mice lethality (Dawes et al., 2019). Nonetheless, we show how some essential human genes may escape capture by known prediction methods since they have 1) no shared model organism phenotype, 2) have different levels of evolutionary conservation, and 3) display variable constraint depending on the mode of inheritance. For instance, at least four genes known to cause a human phenotype do not have a phenotype in mice. To overcome such shortcomings, we show that candidate gene prediction using multiple attributes may also serve to increase sensitivity as overlapping traits appear to enrich for essential genes.

Our findings here support a correlation between essentiality and high embryonic expression. Specifically, embryonic expression is a signature of gene essentiality as it correlates to human and murine essentiality, conservation, and constraint. Highly expressed genes in early embryonic stages, particularly those expressed by the zygote, may play key roles in processes occurring early in development, such as lineage specification, implantation or X-chromosome dosage compensation (Ortega et al., 2018). Highly expressed genes in the human seem to turn on early and occupy essential roles throughout the human lifespan, as they are enriched in basic cellular machinery and metabolic processes, congruent with essential genes in other model systems (Hirsh & Fraser, 2001; Kojima, Tam, & Tam, 2014; Pál, Papp, & Hurst, 2001).

Genome-wide systematic studies of essential genes were pioneered using yeast (Hirsh & Fraser, 2001; Pál et al., 2001). Essential genes to the eukaryotic cell that are absolutely required for the yeast cells to survive also encode core cellular processes (Boone & Andrews, 2015; Fraser, 2015). Yeast essential genes also tend to be conserved across species and highly expressed. More recent genome-wide viability screens in human cells have been performed using CRISPR-Cas9 and gene trap technologies (Fraser, 2015; Hart et al., 2015; T. Wang et al., 2015). In addition, essential genes identified in *in-vitro* studies of haploid human cell lines are helpful. They found that essential genes tended to be enriched in core cellular processes such as translation, transcription and metabolism (Boone & Andrews, 2015). These processes are also highly conserved across species, likely subject to strong negative selection and exhibit high genic constraint (Cardoso-Moreira et al., 2019). Yet translation from *in-vitro* to *in-vivo* may be limited if viability is context dependent (Boone & Andrews, 2015; Fraser, 2015). Future studies that reproduce *in-vivo* biological conditions in early development may help further delineate potential differences. Interestingly, we found that many of our predicted essential genes are important beyond the single cell context, suggesting these genes may be candidates for essentiality at later stages in development.

Our analysis of embryonic gene expression trends suggests that a rise in expression of zygote-activated genes may also be a signature of essentiality. The human embryo undergoes zygotic genome activation at around the 4 to 8-cell stage, a reprogramming process after which development is no longer dependent on oocyte-derived transcripts but is rather driven by *de novo* transcripts (Vassena et al., 2011). In fact, we observed that zygote-activated genes, which had with the highest increase in expression, identified multiple examples of known human essential genes, including *NLRP7* (C. M. Wang et al., 2009). Zygote-activated genes that trend upwards are enriched for essentiality and have a greater tendency to be constrained, suggesting their activation and expression at these stages is critical. Moreover, upregulated genes early in embryonic genome activation vary greatly between species (Xie et al., 2010), further capitulating the importance of direct humans studies.

The aforementioned signatures of essentiality clearly capture overlapping yet distinct sets of essential human genes. We have compiled a list of putative essential gene candidates based on expression data and their relation to other essentiality signatures. These candidates could help with future gene curation, prioritization, and phenotypic correlation analyses, especially as the use of genetic sequencing during pregnancy as a diagnostic tool increases. Moreover, some genes may be uniquely essential in humans, and their identification is instrumental to a deeper understanding of development. Therefore, tools and techniques that identify human-specific essential genes are likely to become increasingly important as we continue to investigate developmental processes that are unique to our species.

